# Early life adversity and borderline intellectual functioning negatively impact limbic system connectivity in childhood: a connectomics-based study

**DOI:** 10.1101/2019.12.13.875161

**Authors:** Valeria Blasi, Alice Pirastru, Monia Cabinio, Sonia Di Tella, Maria Marcella Laganà, Alice Giangiacomo, Gisella Baglio, Michela Zanette, Maria Paola Canevini, Mauro Walder, Mario Clerici, Francesca Baglio

## Abstract

Epigenetic factors related to early life adversity (ELA) in childhood are major risk factors for borderline intellectual functioning (BIF). BIF affects both adaptive and intellectual abilities, commonly leading to school failure and to an increased risk to develop mental and social problems in the adulthood. This study aimed to investigate the neurobiological underpinnings of ELA associated with BIF in terms of global topological organization and structural connectivity and their relation with intellectual functioning.

BIF (N=32) and age-matched typical development (TD, N=14) children were evaluated for intelligence quotient (IQ), behavioral competencies, and ELA. Children underwent an anatomical and diffusion-weighted MR imaging (DWI) protocol. Global brain topological organization was assessed measuring segregation and integration indexes. Moreover, structural matrices, measuring normalized number of fibers (NFn), were compared between the 2 groups using network-based statistics. Finally, a linear regression model was used to explore the relationship between network parameters and clinical measures.

Results showed increased behavioral difficulties and ELA, together with decreased network integration in BIF children. Moreover, significantly lower NFn was observed in the BIF group (p=.039) in a sub-network comprising anterior and posterior cingulate, the pericallosal sulcus, the orbital frontal areas, amygdala, basal ganglia, the accumbens nucleus, and the hippocampus. Linear regression showed that NFn significantly predicted IQ (p<.0001).

This study demonstrated that ELA in children with BIF is associated with a decreased information integration at the global level, and with an altered structural connectivity within the limbic system strictly related to the intellectual functioning.

## 1. Introduction

There is compelling evidence that the epigenetic factors related to early life adversity (ELA) in childhood, such as low socio-economic status (SES), maltreatment, neglect and high levels of parental/family stress, are major risk factors for mental health disorders (1, 2). Moreover, several neuroimaging studies investigating the impact of ELA revealed that low SES, maltreatment and neglect, if experienced during childhood, are associated with abnormal brain function and development in several regions, particularly within the limbic system (3–5). These data have been considered as part of the biological substrate of the “latent vulnerability” (6) according to which the alterations observed at the structural and functional level in several neurobiological systems reflect the (mal)adaptation to neglectful and/or abusive early environments. These changes are likely to be beneficial within the maladaptive context but represent a long-term cost for the subject, thus increasing vulnerability to future stressors (6).

In this context, a particularly vulnerable population is represented by children with borderline intellectual functioning (BIF). BIF is a neuropsychiatric condition characterized by an intelligence quotient (IQ) in the borderline range (70 to 85) associated with adaptive difficulties in social participation (7–9). The major risk factor for the development of BIF is represented by an ELA (10–13). In primary school age, BIF is characterized by limitations in social (14, 15), emotional and behavioral capacities (16, 17). According to recent studies, the prevalence of BIF has been established to be as high as 7 to 12% (7, 18). Children with BIF have a risk as adults to develop mental health problems (e.g. antisocial personality disorder, depression, psychosis, suicide and substance abuse), physical problems and poverty compared to people with average or above average IQ (19–21). Moreover, the presence of BIF negatively impacts the prognosis of all psychiatric diseases (22). Taken together, these data show that BIF is a highly relevant condition for the prognosis and treatment of neuropsychiatric disorders in childhood.

Recently, neuropsychiatric disorders are increasingly being investigated at the level of distributed brain networks rather than in the context of individual brain regions (23, 24). The network-based approach to the study of brain connectivity, so-called brain connectomics, allows the investigation of the topology of the brain as a network. According to graph theory, the brain can be viewed as a complex network constituted of nodes, i.e. the cortical and subcortical gray matter (GM) structures, with pairs of nodes connected by edges, i.e. the white matter (WM) fibers that connect them. This framework allows for the investigation of several properties of the network that define its efficacy and complexity. According to Friston et al (25) there are two fundamental principles of brain organization: functional specialization (segregation) and functional integration according to which complex behaviors derive from the integration of functionally specialized areas that are highly interconnected to form clusters and modules. Graph theory enables the measurement of the metrics exploring this type of organization from a topological point of view.

BIF is a condition characterized by a heterogeneous behavioral phenotype and therefore it is reasonable to think that a distributed network is involved in its clinical manifestations.

The aim of this study was then to investigate the brain network connectivity of children exposed to adverse social environments showing a BIF and its relationship to intellectual functioning. The answer to these questions can have a great impact for the planning of appropriate interventions to prevent the many risks this population faces in the adult age.

We therefore created an ad hoc checklist, the environmental stress check list (ESCL), to assess the adversity that children with BIF were exposed to and compared children with BIF with children with typical development (TD) in terms of brain connectivity and topological organization. Finally, to explore the relationship between intellectual functioning and clinical, environmental and brain connectivity indices, a linear regression approach was used.

## 2. Methods and Materials

### 2.1. Participants

Forty-two children with BIF associated with significant ELA (see later for more details) were recruited from the Child and Adolescent Neuropsychiatry Unit of IRCCS Don Carlo Gnocchi Foundation and the ASST S. Paolo and S. Carlo Hospital.

Inclusion criteria were: 1) age range comprised between 6 to 11 years old; 2) attendance of a primary mainstream school; 3) a Full Scale Intelligence Quotient (FSIQ) score ranging from 70 to 85 determined with the Wechsler Intelligence Scale for Children-III (WISC-III) (26).

A group of eighteen age and sex-matched TD children with a negative history for neurodevelopmental, behavioral or emotional disorders and a FSIQ >85 was also included in the study.

To exclude children with BIF due to biological and/or genetic causes, the presence of any of the following represented an exclusion criteria: 1) major neuropsychiatric disorders (such as ADHD, and autism spectrum disorder); 2) neurological conditions such as epilepsy, traumatic brain injury, brain malformation, infectious disease involving the central nervous system and perinatal complications such as prematurity or other adverse events; 3) systemic diseases such as diabetes or dysimmune disorders, genetic syndromes such as Down syndrome or Fragile X syndrome. Furthermore, a positive history for psychoactive drugs, particularly referring to current or past use of psychostimulants, neuroleptics, antidepressants, benzodiazepines and antiepileptic drugs were also considered exclusion criteria.

All children underwent a neuropsychological evaluation including: the WISC-III (26); the Child Behavioral Checklist (CBCL 6-18) (27); the SES (28); the ESCL, an ad hoc developed check list to explore the environmental stress the children were exposed to (See supplementary Table S1). The ESCL comprised a listing of the V-codes from DSM-5, and Z-codes from ICD-10, exploring Relational, Neglect, Physical, Sexual and/or Psychological Abuse, Educational and Occupational, Housing and Economic, Social Exclusion or Rejection Problems, plus the presence of the following three conditions: social services intervention, major psychiatric diagnosis and/or substance abuse within the family members. The presence of each condition and its relevance for the clinical manifestations was considered and a 0 (absence) to 1 (presence) score was attributed to each item. The ESCL total score could range from 0 to 24. The considered conditions were not weighted for their severity, thus in general higher scores do not represent a more adverse environment.

All subjects underwent a magnetic resonance imaging (MRI) evaluation (see MRI Acquisition section).

The Study was approved by the Ethics Committees of the Don Gnocchi Foundation and of the ASST S. Paolo and S. Carlo Hospital. All parents signed a written informed consent at the first meeting.

### 2.2. MRI Acquisition

MRI was performed on a 1.5 T Siemens Magnetom Avanto (Erlangen, Germany) scanner equipped with a 12-channels head coil. The acquisition included: 1) a 3D T1-weighted Magnetization Prepared Rapid Gradient-Echo (MPRAGE) image, (repetition time (TR)/echo time (TE)=1900/3.37 ms, Filed of View (FoV) = 192×256 mm2, voxel size = 1 mm isotropic, 176 axial slices); 2) a diffusion-weighted (DW) EPI image along 30 directions with b-value=1000 s/mm2 and one without diffusion weighting (TR/TE = 6700/100 ms, FoV = 200×200 mm2, voxel size 1.6×1.6×2.5 mm3, 40 axial slices, two runs); 3) two conventional anatomical sequences (axial PD/T2 and coronal FLAIR) to exclude gross brain abnormalities.

### 2.3. MRI Data analysis

The 3D-T1 images were segmented and parcellated using FreeSurfer version 5.3^1^ into 148 cortical areas (74 for each hemisphere) according to the Destrieux atlas (29). Furthermore, the FreeSurfer automatic labeling process was used to extract seven subcortical regions per hemisphere (thalamus, caudate, putamen, pallidum, and nucleus accumbens, amygdala and hippocampus) and the brain stem for a total of 163 parcels. The quality of recon-all parcellation was assessed in each subject according to ENIGMA guidelines^2^ for cortical and subcortical areas.

Using FMRIB’s Software Library tools^3^, the DW images were corrected for eddy current distortion (30). The motion evaluation was performed by checking the relative movement estimated by eddy toolbox and excluding subjects exciding a threshold fixed to 0.5. Then, using the FSL DTIFIT toolbox^4^ the tensor was estimated for each voxel. The cortical/subcortical parcels were registered to the DW space using the FSL flirt tool (31). Finally, the WM tracts connecting each pair of registered cortical and subcortical parcels (nodes) were reconstructed with TrackVis software^5^.

The connectivity matrices were derived by computing the edges as the number of the reconstructed fibers normalized by the sum of the nodes volumes (NFn) in order to consider the effect of anatomical variability. The matrices were successively employed both for graph and network-based analyses as explained in details in the following sections.

#### 2.3.1. Graph-based analysis

The connectivity matrices were thresholded and binarized. The matrices were thresholded in a way that at least 1/3 of TD children shared the same connections. In order to investigate the topological organization, whole brain network metrics of segregation and integration were derived (32). Specifically, average clustering coefficient (CC), characteristic path length (CPL) and global efficiency (GE) were computed for each subject by means of the Brain Connectivity Toolbox (brain-connectivity-toolbox.net). Furthermore, the density of the matrices was extracted (see Table 1). The resulting indices were compared between the two groups by means of a Mann-Whitney test.

**Table 1.**
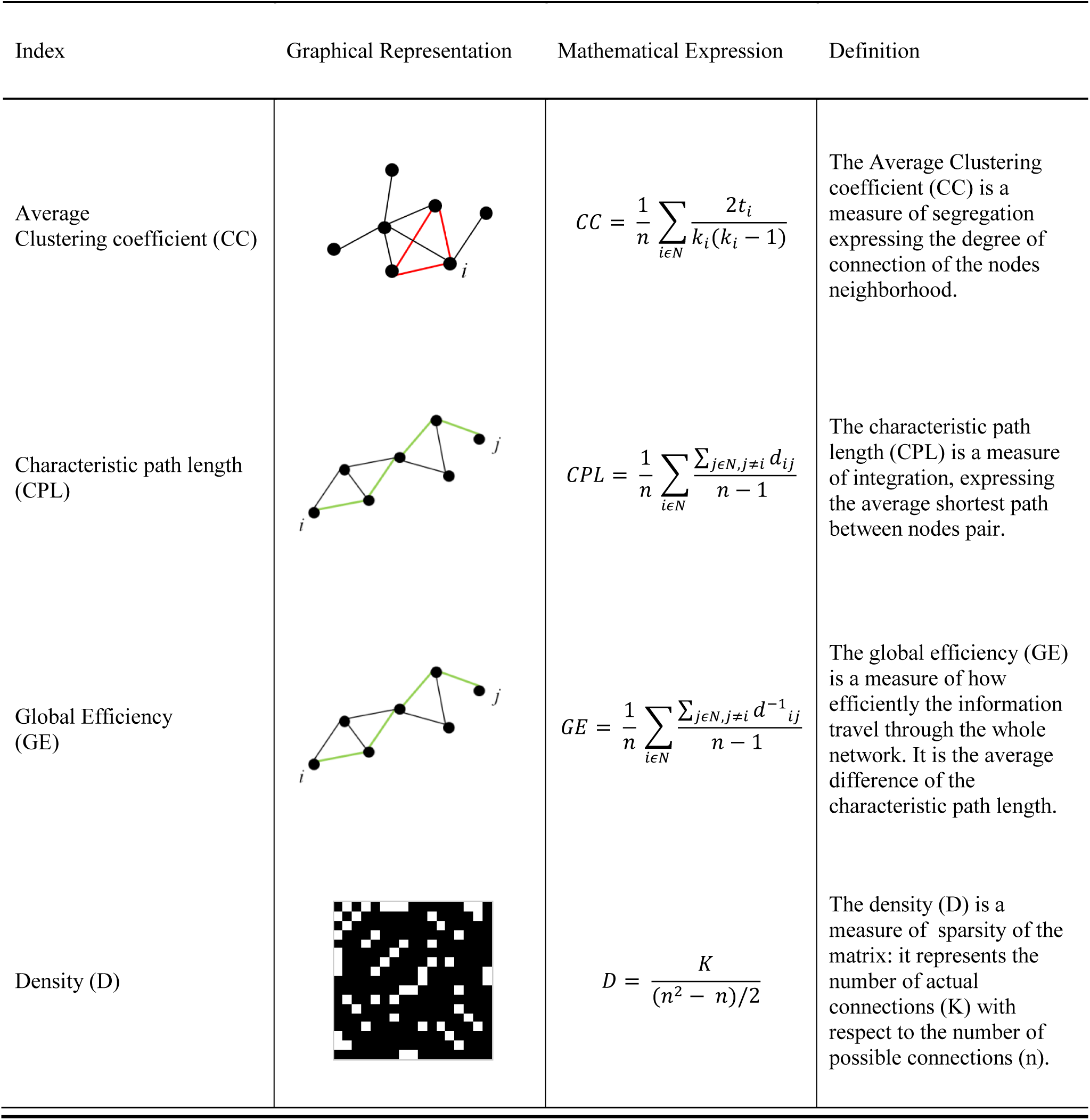
Graph Indices Description. The table shows an overview on the computed global indices (32): specifically for each index a graphical, mathematical and theoretical description is provided. Legend: t=number of triangles of a node neighborhood; d=distance between a pair of nodes; k=number of actual connections (edges different from zero); n=number of possible connections.

| Index | Graphical Representation | Mathematical Expression | Definition |
| --- | --- | --- | --- |
| Average Clustering coefficient (CC) | | $CC = \frac{1}{n} \sum_{i \in N} \frac{2t_i}{k_i(k_i - 1)}$ | The Average Clustering coefficient (CC) is a measure of segregation expressing the degree of connection of the nodes neighborhood. |
| Characteristic path length (CPL) | | $CPL = \frac{1}{n} \sum_{i \in N} \frac{\sum_{j \in N, j \neq i} d_{ij}}{n - 1}$ | The characteristic path length (CPL) is a measure of integration, expressing the average shortest path between nodes pair. |
| Global Efficiency (GE) | | $GE = \frac{1}{n} \sum_{i \in N} \frac{\sum_{j \in N, j \neq i} d_{ij}^{-1}}{n - 1}$ | The global efficiency (GE) is a measure of how efficiently the information travel through the whole network. It is the average difference of the characteristic path length. |
| Density (D) | | $D = \frac{K}{(n^2 - n)/2}$ | The density (D) is a measure of sparsity of the matrix: it represents the number of actual connections (K) with respect to the number of possible connections (n). |

#### 2.3.2. Network-based analysis

Potential group differences were computed using the Network-Based Statistics toolbox (NBS) (33), using an ANCOVA design with age and sex as covariates. The threshold used to identify connections was set to 3.1 (p=.0017), with family-wise error (FWE)-correction using permutation testing (10000 permutations, p=.05). The results were visualized using a circular representations (connectograms,^6^ (34)), according to (35, 36). Finally, to correlate the results deriving from the NBS analysis with the clinical data, the cluster strength (CS) was calculated as the mean strength (weights average) of the sub-network of significant difference between the two groups. A partial correlation (Spearman) with the clinical variables, FSIQ, SES, CBCL, ESCL and the CS, with age and sex as covariates, was then performed. The variables showing a significant correlation were considered as independent variables in a regression model predicting the FSIQ score. All statistical analyses were performed by means of SPSS (Version 25; IBM, Armonk, New York) software.

## 3. Results

### 3.1. Sample and clinical assessment

Due to excessive head movement during the MRI evaluation, 10 children with BIF and 4 TD were excluded from the data analyses, and thus the final sample consisted of thirty-two BIF and fourteen TD children.

Among the 32 children with BIF, 25 had an Adjustment Disorder, 4 had an Anxiety Disorder and 1 had a Disruptive, Impulse-Control, and Conduct Disorder. Moreover, in 14 children a Specific Learning Disorder was associated, in 14 there was a history of Language Development Disorder. Demographic data relative to the 32 BIF and 14 TD children are shown in Table 2. No significant differences were found for age, sex and SES, while the CBCL (p=0.005), the ESCL and as expected the FSIQ (p<0.0001) were significantly different between the two groups (see table S2 for ESCL detailed score).

**Table 2.**
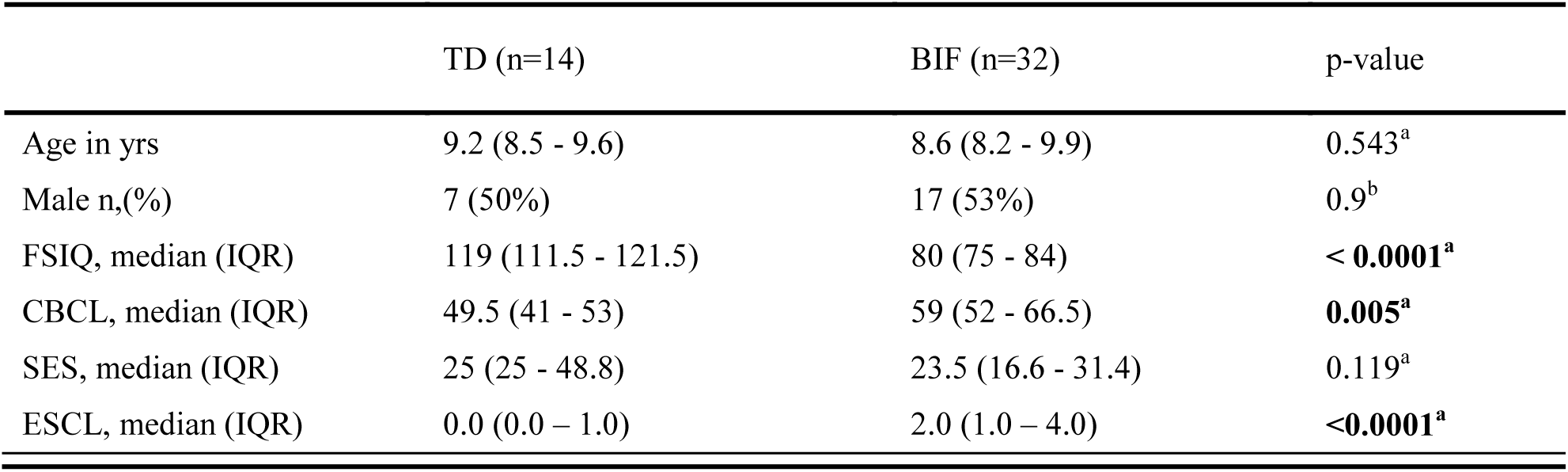
Demographic Variables. IQR=Interquartile range. ^a^Non-parametric (Mann-Whitney test), ^b^Chi-squared test. FSIQ=Full Scale Intelligence Quotient, CBCL=Child Behavior Checklist, SES=Socio-Economic Status, ESCL=Environmental Stress Check List.

### 3.2. Graph-based analysis results

To determine the topological organization of the brain in the two groups, whole brain metrics were derived globally. Specifically, the CC, the CPL and the GE indices were calculated for each subject and compared between the two groups. The density (D) of the matrices was also computed. Table 3 shows results of the Mann Whitney analysis showing significantly increased CPL (p<0.001), and significantly reduced GE (p=0.001) and D (p=0.006) in the group of children with BIF. The CC index, measuring the network segregation, was not different between the two groups.

**Table 3.** Graph-based indexes. Global indices derived from the thresholded and binarized whole brain network matrices for the two groups and their comparison (Non-parametric, Mann-Whitney tests). IQR=Interquartile range.

|  | TD (n=14) | BIF (n=32) | p-value |
| --- | --- | --- | --- |
| Clustering Coefficient, median (IQR) | 0.75 (0.74 - 0.75) | 0.74 (0.74 - 0.76) | 0.252 |
| Characteristic Path Length, median (IQR) | 1.56 (1.54 - 1.56) | 1.58 (1.57 - 1.61) | **0.0002** |
| Global Efficiency, median (IQR) | 0.72 (0.71 - 0.73) | 0.71 (0.70 - 0.72) | **0.001** |
| Density, median (IQR) | 0.69 (0.64 - 0.69) | 0.63 (0.61 - 0.65) | **0.006** |

### 3.3. Network-based analysis results

The NBS analysis comparing the two groups of children in terms of structural connectivity (number of fibers normalized for the volume of the GM parcels) at the network level revealed a sub-network comprising 67 edges and 51 nodes for which the group of children with BIF had a significantly lower NFn compared to the TD group (p=.045 FWE-corrected, see Supplementary Table S3). No significant differences were found in the opposite comparison, TD versus BIF children. The nodes identifying the sub-network comprised the posterior ventral cingulate (vPCC), the striate and extrastriate occipito-temporal cortices, the pericallosal sulcus, the subcallosal gyrus, the hippocampus, the accumbens nucleus and the intraparietal sulcus, bilaterally. The orbital cortex, the insula, the putamen, and the anterior and middle posterior cingulate were involved on the left side while the amygdala, the pallidum, and the superior temporal gyrus (planum polare) on the right side (see Figure 1).

**Figure 1.**
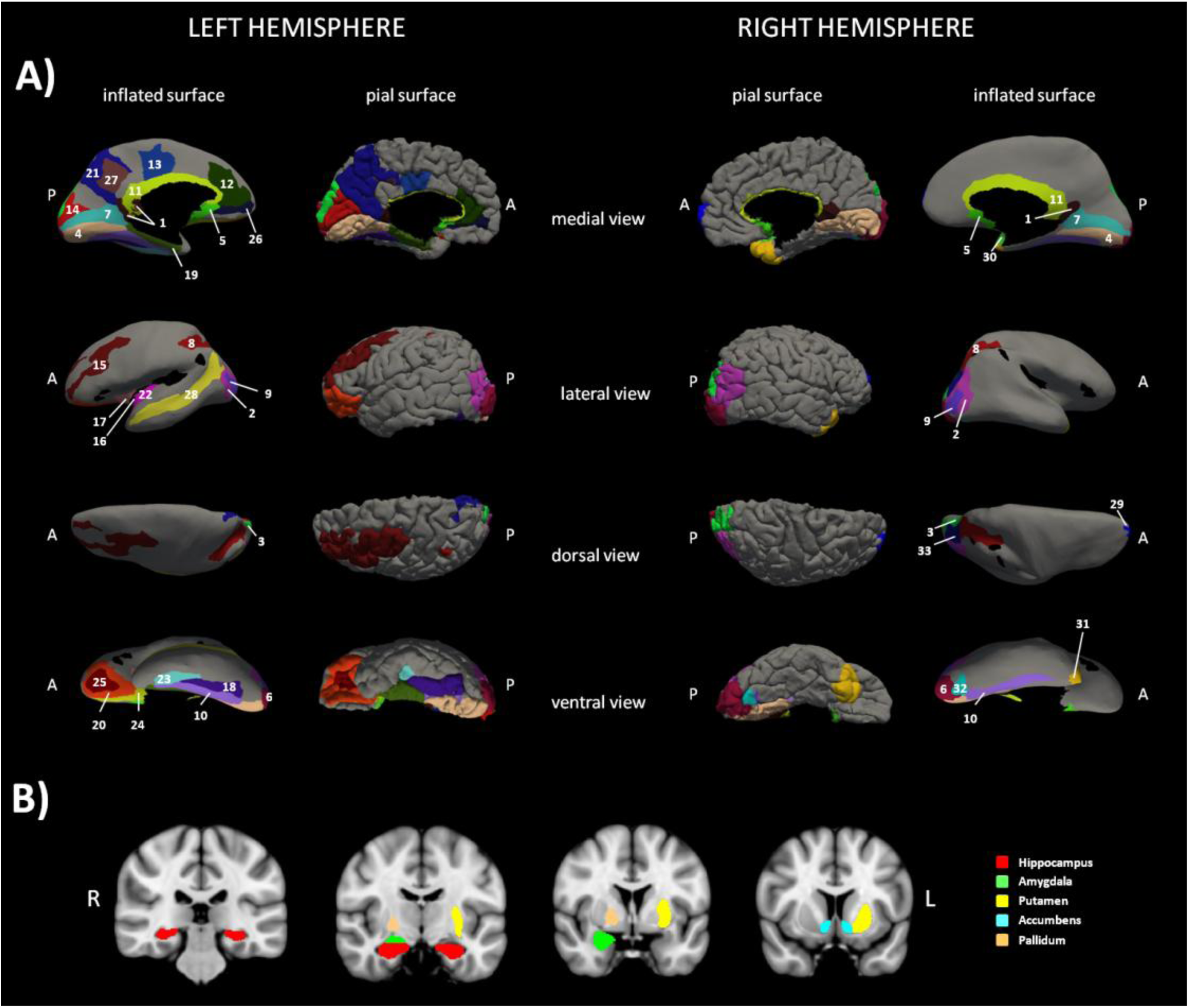
Sub-network cortical/subcortical labels. The figure shows the sub-network of significant difference between the two groups of children obtained with network-based statistic (NBS). Panel A) shows the cortical nodes while panel B) shows basal ganglia belonging to the sub-network. 1=L and R vPCC G, 2=L and R Middle occipital G, 3=L and R Superior occipital G, 4=L and R Lingual part of the medial occipito-temporal G, 5=L and R Subcallosal G, 6=L and R Occipital Pole, 7=L and R Calcarine S, 8= L and R Intraparietal and tansverse parietal S, 9=L and R Middle Occipital and Lunatus S, 10=L and R Collateral and Lingual S, 11=L and R Pericallosal S, 12=L ACC G and S, 13=L pMCC G and S. 14=L Cuneus G, 15=L Middle Frontal G, 16= L Long Insular G and central S of the Insula, 17=L Short Insular G, 18=L Fusiform G, 19=L Parahippocampal part of the medial occipito-temporal G, 20=L Orbital G, 21=Precuneus G, 22= L Inferior Segment of the Circular S of the Insula, 23=L Anterior Transverse Collateral S, 24=L Medial Orbital (Olfactory) S, 25= L Orbital (H Shaped) S, 26=L Suborbital S, 27=L Subparietal S, 28=L Superior Temporal S, 29=R Transverse Frontopolar G and S, 30=R Planum polare of the Superior Temporal G, 31=R Temporal Pole, 32=R Posterior Transverse Collateral S, 33= R Superior occipital and Transverse Occipital S. G=Gyrus/i, S=Sulcus/i; R=right hemisphere, L=left hemisphere, ACC=anterior cingulate cortex, pMCC=middle posterior cingulate cortex, vPCC=ventral part of the posterior cingulate cortex.

In Figure 2 the 67 edges connecting the nodes of the identified sub-network are represented on a connectogram (see Supplementary Table S3 for a complete listing).

**Figure 2.**
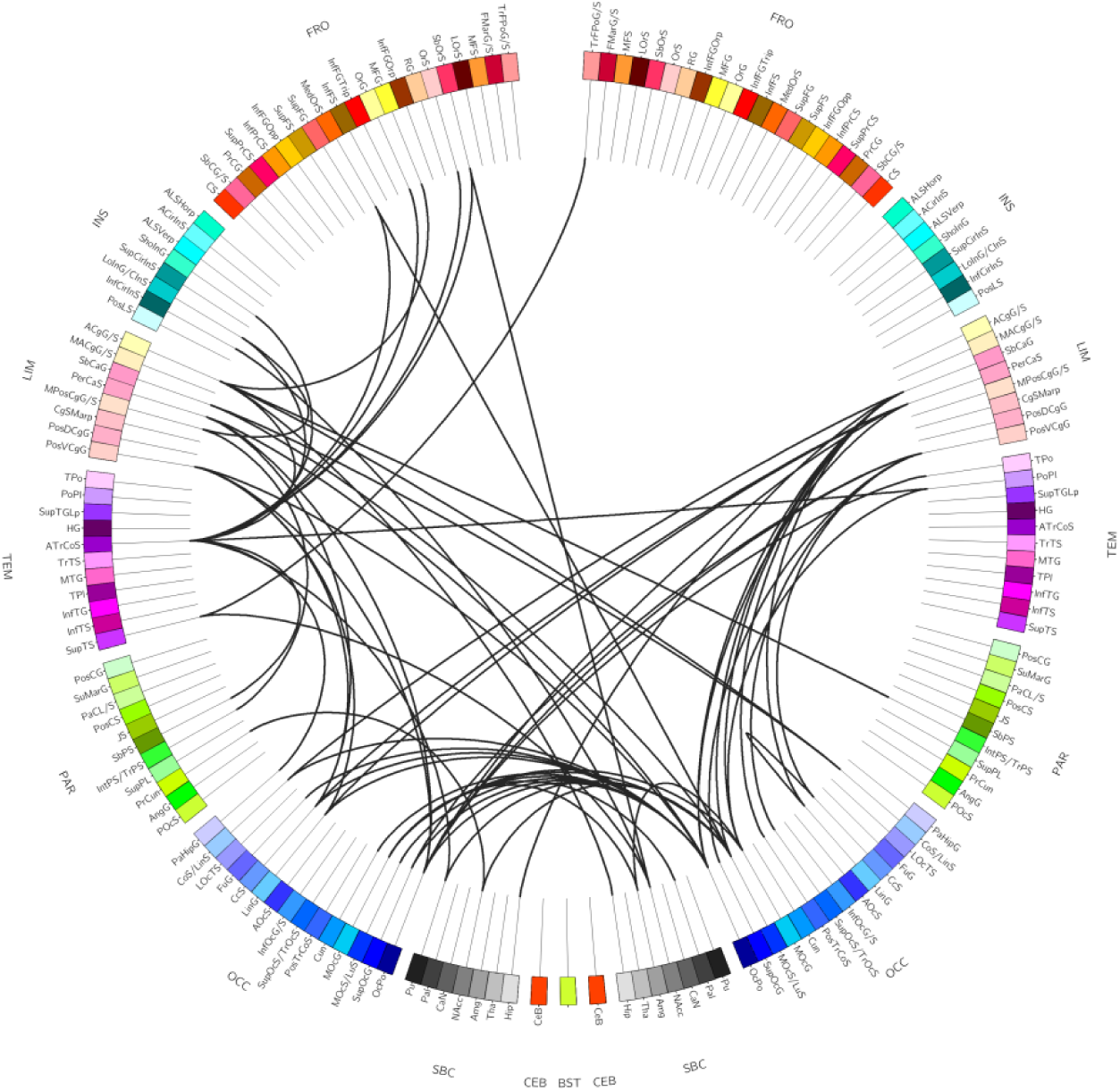
Connectogram-based edges representation. The figure shows the circular representation of the edges belonging to the sub-network of significant difference derived from network-based statistic (NBS). For a complete listing of the nodes involved see Supplementary Table 2.

The partial correlation analysis between the clinical variables FSIQ, SES CBCL total score, ESCL and the CS, with age and sex as covariates, resulted significant for the FSIQ and the CS (r= 0.71; p<0.0001), FSIQ and CBCL (r=-0.47; p=0.003) and FSIQ and ESCL (r=-0.45, p=0.004). The correlation between FSIQ and SES did not survive statistical threshold (r=0.24, p=0.144) According to the correlation results, the linear regression model comprised FSIQ as the dependent variable and CS, CBCL and ESCL as independent variables. Results revealed that only the CS had a significant predictive value for the FSIQ (R2=0.58;beta=0.6; t=5.0; p<0.0001).

## 4. Discussion

Our study focused on the neurobiological underpinnings of ELA associated with BIF. To this purpose we investigated brain network topological organization and connectivity in 32 children with BIF and 14 age -matched TD children. All children were clinically characterized for exposure to environmental stress, intellectual functioning and behavioral characteristics.

The interest in children with BIF condition is in line with the emerging concept of a preventative psychiatry oriented towards the development early intervention models for mental health making it a priority to intervene in the developmental period. This approach can benefit from the understanding of the neurocognitive mechanisms involved in psychiatric disorders that share common ethiopatogenetic mechanisms rather than conventional diagnostic boundaries (37). Children with BIF indeed represent a population highly vulnerable to the development of major psychiatric conditions as adults (8, 20, 22) with environment-related etiopathogenetic factors (10–13).

The results of this study showed that the children with BIF were characterized by significantly more environmental stressful elements and behavioral difficulties compared to the TD children. The major sources of stress in the group of children with BIF were related to educational problems, family disaggregation and /or the intervention of social services for impossibility of the parents to cope. Moreover, in few cases there was a history of parents’ drug abuse, or abandonment of children by the father, or father in prison. Some children grew up away from their parents. In few cases, the low SES resulted in inadequate housing in a way that was considered relevant to the diagnosis. This is in line with data from recent studies, one of which investigated a very large cohort (14,000 children) and showed that children from low SES families scored on average 6 IQ points lower at age 2 than children from high SES families; by age 16, this difference had almost tripled (38). Moreover, it has been demonstrated that low SES affects both learning abilities and brain development in regions critical for memory and emotion regulation such as the hippocampus and the amygdala (3). In our study, the SES was not statistically different between the two groups of children but the BIF group was exposed to significantly greater environmental stress. Since previous neuroimaging studies have demonstrated that maltreatment and/or neglect in childhood can impact brain development (4, 39–41) our study investigated the neural bases of the BIF condition. In a previous study from our group, abnormal brain development of the parahippocampal, temporal and sensory-motor cortices in children with BIF was found with a single brain region approach (42). In this study, we used a brain connectomics approach, and the whole brain topological organization was investigated to capture indices of global organization by using a graph-based analysis. Results showed significant differences between the two groups in the CPL and in the GE. Both indices explore the level of integration of the information coming from different brain structures and the observed differences indicate lower levels of integration in the group of children with BIF. The lower level of integration reasonably reflects the broad range of cognitive and behavioral difficulties observed in children with BIF ranging from specific learning disorder to difficulties in higher order functions such as executive functions, planning, inhibition, attention and behavior (7). These data are in line with a study on healthy adult subjects showing that higher intelligence scores corresponded to a shorter CPL and a higher GE of the networks, indicating a more efficient parallel information transfer in the brain (43).

Beside the graph-based analyses, we further investigated structural connectivity using a network-based approach. Results revealed significant between-group differences in a sub-network connecting several cortical and subcortical areas, mostly related to the limbic system. Specifically, the anterior and posterior cingulate cortices, the hippocampus and the parahippocampus, the pallidum, the putamen, and the accumbens nucleus, the subcallosal and pericallosal cortices, the frontal-orbital regions, and the extra-striate visual cortices were part of the network. To our knowledge this is the first report of the involvement of the limbic system connectivity in children exposed to ELA showing a BIF and is in line with consistent data from neuroimaging studies showing that abuse, maltreatment, and neglect in childhood are associated with specific epigenetic and neural signatures related to long lasting structural and functional changes in brain areas belonging to the limbic system (4, 37, 44) together with alteration in the structural connectivity at the network level (39). In particular, it has been shown that adults who experienced maltreatment during childhood show hyper-responsiveness of the amygdala to fearful stimuli (45, 46), even during pre-attentive conditions (47, 48), hypo-reactivity of the hippocampus to pleasant autobiographical stimuli and hyper-reactivity during unpleasant stimuli (47), and abnormal reactivity to reward in the nucleus accumbens (49). Moreover, at the morphometric level reduced cortical thickness in the anterior cingulate, superior frontal gyrus, and orbitofrontal cortex, reduced cortical surface area in the left middle temporal area and lingual gyrus, and gyrification deficits in the lingual gyrus and the insula were demonstrated (4). Finally, altered structural brain network topology was found at the global and lobar level in maltreated children with normal IQ, with significant reductions of the connectivity strength and increment in the CPL, both related to neural integration capacity (39). These data thus showed functional and structural alterations in the neurocognitive systems involved in fear – emotion regulation, motivation – reward, and learning processes, which have been considered a sort of early developed adaptive calibration of these systems to adverse environments. In turn, these changes may represent a “latent vulnerability” to future stressors associated with an increased risk of developing mental health disorders later in life (6).

In the present study the clinical population involved children exposed to several environmental stressors associated with BIF in the presence of clinical manifestations ranging from neurodevelopmental disorders such as specific learning disorder, language and movement development disorders to adjustment or behavioral or anxiety disorders. Therefore, BIF cannot be considered a latent condition but a clinically manifest one that necessitates immediate interventions. Moreover, the involvement of the whole limbic system is in line with the clinical manifestations of these children ranging from the emotion regulation/behavioral problems, difficulty in inhibiting impulsive responses, and the motivational problems (9) and with the poor prognosis in terms of risk to develop psychiatric disorders in the adult age (19–21). The limbic system has a pivotal role in all these aspects (50).

To investigate the causal relationship between the IQ and the clinical, neural, and environmental aspects a linear regression approach was used. Results showed that the cluster strength of the altered network was a predictor of the IQ of the children, while the clinical (CBCL) and environmental (ESCL) variables were not. These data demonstrate a strict relationship only between the network connectivity and the intellectual functioning of our children. This could be due either to a non linear relation between the other variables or to the absence of such a relationship. Moreover, the ESCL was not a weighted measure and thus did not reflect the severity of the environmental condition.

Despite the great innovativeness of the present study in shedding new light on the pathways associated with BIF, our study is not free from some limitations. In particular, the number of diffusion directions in the DWI sequence might appear as limited when compared to the state-of-the-art MRI acquisitions. However, considering the particular cohort of subjects included, more prone to movement during the examination (14 DWI datasets were discarded for elevated head motion), the chosen DTI sequence represents a good trade-off between having qualitatively good data and acquisition time.

Taken together all these data indicate that the neurobiological underpinning of the clinical manifestations of children exposed to ELA associated with BIF is represented by the reduced GE in information integration and by the altered structural connectivity in the circuitry crucial for the regulation of emotions, behavior, motivation, and memory. These abnormalities are closely related to the IQ of children. We consider these data extremely relevant for the understanding of the cognitive and behavioral manifestations of these children and for the implementation of appropriate interventions able to reduce the risk of future psychiatric disorders in these children.

## 5. Acknowledgments

The authors wish to thank Dr. Niels Bergsland for his careful revision of the English language, and all the children and their families for participating in the study.

## 6. Author Contributions Statements

All authors participated in writing and critically reviewing the work, providing important intellectual content and approving the final form.

VB, MAC and FB designed and supervised the research, interpreted the results and drafted the manuscript. AP and MOC performed MRI data analysis and drafted the manuscript. SD and MML performed the statistical analysis. MZ, MPC and MW recruited patients, AG and GB performed the behavioral and neuropsychological evaluations; MZ also contributed in the analysis of the clinical data.

## Supplementary materials

**Supplementary Table S1.** Environmental Stress Check List.

| DSM V V-code<br>(ICD-10 Z-code) | Environmental stressful condition and/or problems | Score<br>(1/0)* |
| --- | --- | --- |
| V61.8 (Z62.891) | Sibling Relational problem |  |
| V61.8 (Z62.29) | Upbringing away from parents |  |
| V61.8 (Z63.8) | High expressed emotion level within family |  |
| V61.20<br>(Z62.820) | Parent-child relational problem |  |
| V61.29<br>(Z62.898) | Child affected by parental relationship distress |  |
| V61.03 (Z63.5) | Disruption of family by separation or divorce |  |
| (Z63.2) | Inadequate family support |  |
| V61.21<br>(Z69.010) | Encounter for mental health services for victim of parental child physical abuse |  |
| V61.21<br>(Z69.020) | Encounter for mental health services for victim of non-parental child physical abuse |  |
| V61.21<br>(Z69.010) | Encounter for mental health services for victim of parental child sexual abuse |  |
| V61.21<br>(Z69.020) | Encounter for mental health services for victim of non-parental child sexual abuse |  |
| V61.21<br>(Z69.010) | Encounter for mental health services for victim of parental child neglect |  |
| V61.21<br>(Z69.020) | Encounter for mental health services for victim of non-parental child neglect |  |
| V61.21<br>(Z69.010) | Encounter for mental health services for victim of parental child psychological abuse |  |
| V61.21<br>(Z69.020) | Encounter for mental health services for victim of non-parental child psychological abuse |  |
| V62.3 (Z55.9) | Academic or educational problem (Underachievement in school) |  |
| V62.3 (Z55.9) | Academic or educational problem (School-Family conflicts) |  |
| V60.1 (Z59.1) | Inadequate Housing |  |
| V60.2 (Z59.6) | Low income |  |
| V62.4 (Z60.3) | Acculturation difficulty |  |
| V62.4 (Z60.4) | Social exclusion or rejection |  |
|  | Social Services Intervention |  |
|  | Major Psychiatric Diagnosis within the family |  |
|  | Substance abuse within the family |  |

**Supplementary Table S2.** Environmental Stress Scoring. Prevalence of Environmental Stress factors in the group of children with Borderline Intellectual Functioning (BIF) and with Typical Development (TD).

|  | BIF<br>group | TD<br>group |
| --- | --- | --- |
| V60.1 (Z59.1) - Inadequate Housing | 4 | - |
| V60.2 (Z59.6) - Low income | 6 | - |
| V61.03 (Z63.5) - Disruption of family by separation or divorce | 10 | 1 |
| V61.20 (Z62.820) - Parent-child relational problem | 6 | - |
| V61.21 (Z69.010) Encounter for mental health services for victim of parental child physical abuse | - | - |
| V61.21 (Z69.020) - Encounter for mental health services for victim of non-parental child physical abuse | - | - |
| V61.21 (Z69.010) - Encounter for mental health services for victim of parental child sexual abuse | - | - |
| V61.21 (Z69.020) - Encounter for mental health services for victim of non-parental child sexual abuse | - | - |
| V61.21 (Z69.010) Encounter for mental health services for victim of parental child neglect | 4 | - |
| V61.21 (Z69.020) - Encounter for mental health services for victim of non-parental child neglect | - | - |
| V61.21 (Z69.010) - Encounter for mental health services for victim of parental child psychological abuse | 1 | - |
| V61.21 (Z69.020) - Encounter for mental health services for victim of non-parental child psychological abuse | - | - |
| V61.29 (Z62.898) - Child affected by parental relationship distress | 6 | - |
| 1.V61.8 (Z62.891) - Sibling Relational problem | 0 | - |
| V61.8 (Z62.29) - Upbringing away from parents | 3 | - |
| V61.8 (Z63.8) - High expressed emotion level within family | 5 | 3 |
| V62.3 (Z55.9) - Academic or educational problem (Underachievement in school) | 29 | - |
| V62.3 (Z55.9) - Academic or educational problem (School-Family conflicts) | 10 | - |
| V62.4 (Z60.3) - Acculturation difficulty | 3 | - |
| V62.4 (Z60.4) - Social exclusion or rejection | 1 | - |
| (Z63.2) - Inadequate family support | 5 | - |
| Social Services Intervention | 5 | - |
| Major Psychiatric Diagnosis within the family | 2 | - |
| Substance abuse within the family | 2 | - |

**Supplementary Table S3.** Edges composing the network of statistical differences. The sub-network presents the following characteristic: number of nodes=51; number of edges=67; p-value=0.045. L=left hemisphere; R=right hemisphere; S=Sulcus/i; G=Gyrus/i; ACC=Anterior Cingulate Cortex; pMCC=Middle-posterior Cingulate Cortex; vPCC=Posterior-ventral part of the Cingulate Cortex. The parcels labeling is the one reported in (1).

| Sub-Network Edges (n=67) and t-values |  |  |
| --- | --- | --- |
| L Anterior Transverse Collateral S | L Subparietal S | 3.1 |
| R Subcallosal G | R Lingual part of the medial occipito-temporal G | 3.11 |
| L Cuneus | R Occipital Pole | 3.11 |
| L Subcallosal G | R Middle Occipital and Lunatus S | 3.12 |
| L Superior Temporal S | R Accumbens | 3.12 |
| L Superior Occipital G | R vPCC G | 3.13 |
| L Occipital Pole | R Temporal Pole | 3.13 |
| R vPCC G | R Calcarine S | 3.14 |
| L Accumbens | R Middle Occipital and Lunatus S | 3.14 |
| L Fusiform G | R Occipital Pole | 3.14 |
| L Middle Occipital G | R Amygdala | 3.14 |
| L Superior Temporal S | R Transverse Frontopolar G and S | 3.15 |
| L Collateral and Lingual S | R Subcallosal G | 3.15 |
| L Subcallosal G | R Collateral and Lingual S | 3.16 |
| L ACC G and S | L Anterior Transverse Collateral S | 3.17 |
| L Suborbital S | L Anterior Transverse Collateral S | 3.19 |
| L Putamen | R Middle Occipital and Lunatus S | 3.19 |
| R Middle Occipital G | R Amygdala | 3.19 |
| L Lingual part of the medial occipito-temporal G | R Occipital Pole | 3.2 |
| L pMCC G and S | L Intraparietal and transverse parietal S | 3.23 |
| R vPCC G | R Superior occipital and Transverse Occipital S | 3.23 |
| L Long Insular G and central S of the Insula | L pMCC G and S | 3.24 |
| L Superior Occipital G | R Occipital Pole | 3.24 |
| R Calcarine S | R Lingual part of the medial occipito-temporal G | 3.29 |
| L Inferior Segment of the Circular S of the Insula | R Accumbens | 3.31 |
| L Collateral and Lingual S | R Occipital Pole | 3.32 |
| L Hippocampus | R Subcallosal G | 3.33 |
| L Anterior Transverse Collateral S | R Planum polare of the Superior Temporal G | 3.33 |
| L Occipital Pole | R vPCC G | 3.38 |
| L Middle Occipital G | R Pallidum | 3.38 |
| R Subcallosal G | R Middle Occipital G | 3.4 |
| L Middle Occipital G | R Middle Occipital and Lunatus S | 3.42 |
| L Anterior Transverse Collateral S | L Parahippocampal part of the medial occipito-temporal G | 3.43 |
| L vPCC G | L Occipital Pole | 3.43 |
| R Subcallosal G | R Posterior Transverse Collateral S | 3.48 |
| L Suborbital S | R Accumbens | 3.51 |
| L ACC G and S | R Intraparietal and transverse parietal S | 3.52 |
| L Long Insular G and central S of the Insula | L Anterior Transverse Collateral S | 3.59 |
| L Middle Frontal G | L ACC G and S | 3.6 |
| L Middle Occipital G | R Middle Occipital G | 3.6 |
| L Lingual part of the medial occipito-temporal G | R Pericallosal S | 3.61 |
| R Planum polare of the Superior Temporal G | R Middle Occipital G | 3.65 |
| L ACC G and S | R Collateral and Lingual S | 3.66 |
| R vPCC G | R Superior Occipital G | 3.68 |
| L ACC G and S | R Hippocampus | 3.68 |
| R Subcallosal G | R Middle Occipital and Lunatus S | 3.75 |
| L Pericallosal S | L Calcarine S | 3.76 |
| R Subcallosal G | R Occipital Pole | 3.76 |
| L Medial Orbital (Olfactory) S | L Anterior Transverse Collateral S | 3.82 |
| L Medial Orbital (Olfactory) S | R Middle Occipital and Lunatus S | 3.85 |
| L Precuneus | L Accumbens | 3.86 |
| L vPCC G | R Occipital Pole | 4.06 |
| L Middle Occipital and Lunatus S | R Occipital Pole | 4.09 |
| L Anterior Transverse Collateral S | L Putamen | 4.12 |
| L vPCC G | L Lingual part of the medial occipito-temporal G | 4.21 |
| L Orbital (H Shaped) S | L Anterior Transverse Collateral S | 4.3 |
| L Short Insular G | L Anterior Transverse Collateral S | 4.31 |
| L vPCC G | L Calcarine S | 4.32 |
| L Pericallosal S | R Occipital Pole | 4.38 |
| L Middle Occipital G | R Occipital Pole | 4.43 |
| L Pericallosal S | L Occipital Pole | 4.49 |
| L Orbital G | L Anterior Transverse Collateral S | 4.7 |
| L Calcarine S | R Occipital Pole | 4.7 |
| L Calcarine S | R Pericallosal S | 4.81 |
| L Occipital Pole | R Occipital Pole | 4.81 |
| L Occipital Pole | R Pericallosal S | 6.07 |
| R Pericallosal S | R Occipital Pole | 6.3 |

## Footnotes

1 https://surfer.nmr.mgh.harvard.edu/

2 http://enigma.ini.usc.edu/enigma-vis/

3 FSL; http://www.fmrib.ox.ac.uk/fsl

4 https://fsl.fmrib.ox.ac.uk/fsl/fslwiki/FDT

5 http://trackvis.org

6 http://circos.ca/

